# Loss of dispersal via ice nucleation activity constrains microbial evolution

**DOI:** 10.64898/2026.02.11.704803

**Authors:** Jonathan M. Jacobs, Zachary Konkel, Jelmer W. Poelstra, Jules Butchachas, Lillian Ebeling-Koning, Galit Renzer, Konrad Meister, Jason C. Slot, Stephen P. Cohen, Verónica Roman-Reyna, Cindy E. Morris

**Author notes:** Corresponding authors: Jonathan M. Jacobs, Cindy E. Morris. **Author Contributions:** ZK, JP, JB, CM, JJ conceptualized the research; ZK, JP, JB, JJ, SPC, JS, VRR carried out the research and investigations. Visualizations were created by ZK, JP, SPC, VRR. JJ, VRR, CM acquired funding for this project. JJ and CM coordinated and supervised this project. JJ and CM. ZK, JP, JB, CM, JJ, SPC, VRR, LEK, JS participated in writing and editing the manuscript. **Competing Interest Statement:** Authors declare that they have no competing interests.

## Abstract

The ability to disperse over long distances through the atmosphere is a common trait across the tree of life, facilitating resource access and increasing long-range gene flow. Loss of dispersal mechanisms, *viz*. flight, can occur in animals found on islands where documented phenotypic changes like loss of wingspan impedes longer distance travel to mate with the metapopulation. Bacteria also experience atmospheric flight and descend via bioprecipitation by catalyzing the freezing of cloud droplets with protein InaZ. InaZ triggers ice nucleation at temperatures near 0_∘_C(1). This ice nucleation activity (INA), a biophysical trait, enhances bacterial deposition through precipitation. The role of InaZ-mediated ice nucleation on bacterial dispersal is well documented, but the impact of loss of INA and thus reduction or loss of atmospheric dispersal on bacterial ecology and evolution has not been described. Here we show that the loss of the ancestral *inaZ* gene restricts bacterial dispersal and leads to significant genetic and ecological isolation across multiple genera. Through the analysis of available complete genomes, we demonstrate that lineages lacking functional *inaZ* experience major gene loss events, reduced recombination rates and a marked dependence on human-mediated or insect transmission. These INA-lacking bacteria exhibit an increased ecological signature of isolation that parallels the distribution of geographically isolated animals. Our results establish InaZ as a keystone biophysical trait that defines microbial dispersal strategies. We anticipate these findings will provide a framework for understanding how shifts in biophysical traits drive niche differentiation and changes in dispersal with downstream consequences for Earth system processes.

**Significance Statement:** Some microorganisms catalyze freezing of cloud droplets near 0°C via ice nucleation activity (INA) enhancing their deposition. We determined that loss of the gene encoding the INA protein in Gammaproteobacteria restricts bacterial dispersal. Bacteria that lost this ancestral trait compared to relatives with INA experienced distinct, major gene loss events, altered gene flow and marked dependence on transmission by plant tissues or insects and an increased ecological signature of isolation paralleling the geographically isolated plants and animals. We posit that gene loss for biophysical traits such as INA is a keystone example of the consequences of a biological trait defining microbial dispersal.

## Introduction

The ability to disperse over long distances through the atmosphere is a common trait found across the tree of life (2). Plants and fungi have adaptations that facilitate wind-borne dispersal of seeds, spores and gametes, whereas various invertebrate and vertebrates are capable of gliding and active flight (2–4). Wind-borne movement and flight expand resource access and increase long-range gene flow for genetic exchange (3). The capacity for air-borne movement can be lost or impeded. For example, birds can be isolated on islands or lose the capacity to fly. This is associated with the emergence of distinct phenotypes such as wing length reductions or genome-wide deleterious mutations due to the resulting genetic isolation (5, 6).

Microorganisms that spread aerially depend on passive means for long distance airborne movement. This involves emissions and entrainment out of the planetary boundary layer, movement with air masses in the free troposphere and deposition by settling or scrubbing by rain and snowfall (7). Whereas meteorological phenomena and atmospheric physics reign most of the underlying mechanisms, microorganisms with ice nucleation activity (INA) enrich their deposition by initiating the freezing of cloud droplets, a step crucial for precipitation in temperate climates (8). In bacteria, deposition with precipitation - and thus effective dispersal - is facilitated by the InaZ protein, which promotes water to form ice at temperatures much warmer (near -2° C, for example) than those of other natural ice nucleators (9, 10). This highly conserved protein was initially described as being confined to the Gammaproteobacteria (11). Our analysis of whole genomes in the NCBI database confirmed that InaZ is found in three environmental Gammaproteobacteria genera: *Pantoea, Pseudomonas*, and *Xanthomonas*, which is consistent with isolations from precipitation (Fig. S1). However *inaZ* presence was inconsistent (Fig. 1) across the three genera leading us to explore the evolutionary drivers for INA. Furthermore, we examined if changes in INA led to genomic signatures typical of organisms with impaired capacity for dispersal.

**Figure 1.**
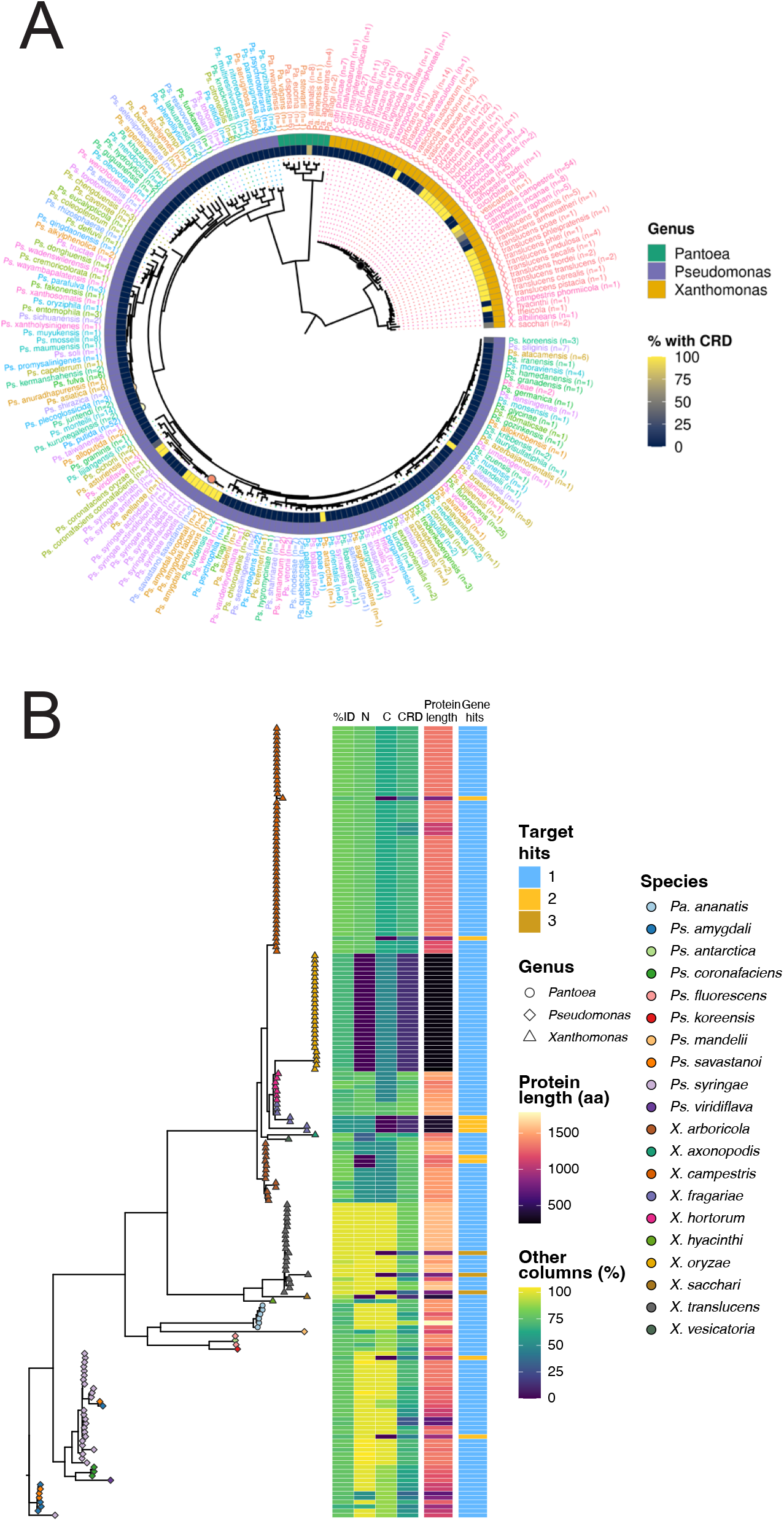
InaZ is ancestral in Gammaproteobacteria with nonuniform distribution. A maximum-likelihood phylogenomic tree was created from (A) one representative bacterial genome per species or (B) 231 InaZ identified homologs. (A) InaZ was considered absent (dark blue) or present (yellow) across *Pseudomonas, Xanthomonas* and *Pantoea*. (B) We created a novel *in silico* approach that predicts InaZ function from the three functional domains: an N-terminus, central repeat domains (CRDs) that comprise predictable repeat sequences, and a C-terminus (15, 16).

## Results and Discussion

### *inaZ* is ancestral and vertically-inherited

Gammaproteobacterial *Pantoea, Pseudomonas*, and *Xanthomonas* are often associated with plant disease and can elicit frost damage in an InaZ-dependent manner (*9*). Gammaproteobacteria collectively are important leaf community members, and their genome evolution has been driven by plant colonization (12). Much of the research on *Pantoea, Pseudomonas*, and *Xanthomonas* evolution has been through the lens of plant leaf colonization and pathogenesis, even though INA tightly links these bacteria to the water cycle (8, 10, 13, 14).

To determine the role of ice nucleation in bacterial genome evolution, we first examined the inheritance of *inaZ* across the Gammaproteobacteria, the only group of bacteria where InaZ is found (11). We reconstructed a maximum-likelihood core gene phylogenomic tree derived from 1531 *Pantoea, Pseudomonas*, and *Xanthomonas* complete genomes, as well as a protein tree of 231 InaZ identified homologs. The phylogenomic species tree of *inaZ-*containing Gammaproteobacteria was generally consistent with the *InaZ* gene tree (Fig. 1; Fig. S2).

The most parsimonious explanation for this topological congruence is that *inaZ* has been vertically inherited in Gammaproteobacteria. This is further supported by colocalization of *InaZ* with flanking genes in each genus and by intragenus synteny conservation of the *inaZ* gene cluster (Fig. 2). Across genera, however, we determined that InaZ is retained without conserved, colocalized adjacent homologs. We also identified a possible rare case of horizontal transfer in *Pseudomonas*, including in two *P. fluorescens inaZ* strains, which are placed in a position that deviates from the species tree with high confidence (Fig. S2).

**Figure 2.**
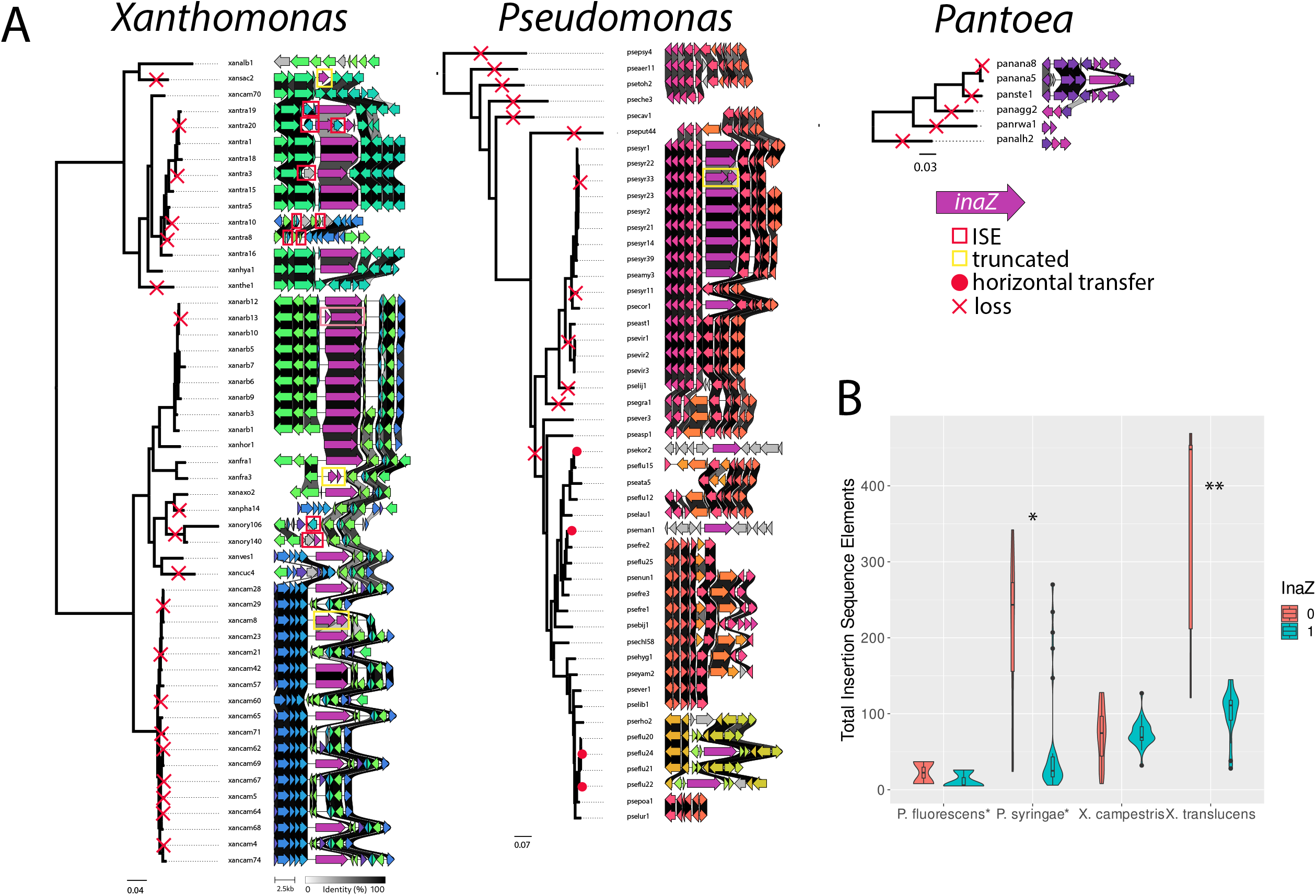
Pathways toward *inaZ* loss and association with transposable element expansion. A) Maximum likelihood phylogenomic trees created from single copy orthologs from *Pseudomonas, Xanthomonas* or *Pantoea* with corresponding *inaZ* clusters. Loss is depicted by a red X. Red dots signify horizontal transfer events. Boxes signify a transposon-mediate loss (red) or a fragmented *inaZ* (yellow). B) ISE abundance calculated in comparison to *inaZ* presence or absence across *Pseudomonas* and *Xanthomonas* species.

### *inaZ* is frequently lost in Gammaproteobacteria

In light of the numerous strains that lacked *inaZ*, we aimed to determine if these effectively represent gene loss events. Among the strains characterized, we observed *inaZ* mutation events, and therefore aimed to define the mechanisms of loss. We found clusters where *inaZ* was completely absent or inactivated by truncation and/or fragmentation, which sometimes was disrupted by a transposable insertion sequence element (ISE) (Fig. 2). This is consistent with the nonuniform occurrence of the gene across the species tree (Fig. 1).

ISEs were frequent causal agents of *inaZ* mutation and inactivation (Fig. 2). ISEs were recovered in loci that contain or now lack *inaZ* from *X. translucens, P. syringae, X. campestris*, and *P. fluorescens*. We identified ISEs truncating, adjacent to or in place of loci that are assumed to have contained *inaZ* ancestrally in 10 representative lineages (Fig. 2). We randomly selected genomes in each clade of *inaZ-*containing and lacking genomes and identified ISEs as causal agents of their loss when they were observed truncating, directly abutting or positioned in *inaZ*. In addition to ISE-mediated loss, we also observed a subset of frameshift mutations resulting in vestigial *inaZ* pseudogenes in *Xanthomonas* and *Pseudomonas* (Fig. 2).

We found that ISE expansion is a mechanism of *inaZ* loss in multiple lineages. *inaZ* mutation in *P. syringae* lineages correlated with ISE expansion (Welch’s t-test p < 0.001), but we were not able to exclude cocorrelation with phylogenetic relatedness in this clade (p = 0.174) (Fig. 3B). The average ISEs present in *X. campestris* were not distinct for genomes lacking or containing *inaZ*, (Fig. 2B). We frequently observed that general ISE expansion is associated with *inaZ* mutations in *X. translucens* (Fig. 3B), which remains statistically significant when controlling for phylogenetic relatedness (p = 0.006). There were multiple examples of independent disruptions of *inaZ* by transposable elements including IS*110*, IS*3*, IS*4* and IS*1595*, suggesting a common mechanism for loss in this species (Fig. 2A, Fig. S4). This frequent *inaZ* loss is consistent with our observation of INA in *X. translucens* in our culture collection (n= 284 strains). 252 strains representing four continents were INA and froze water at –6°C; while 32 were ice nucleation inactive (Fig. S3).

**Figure 3.**
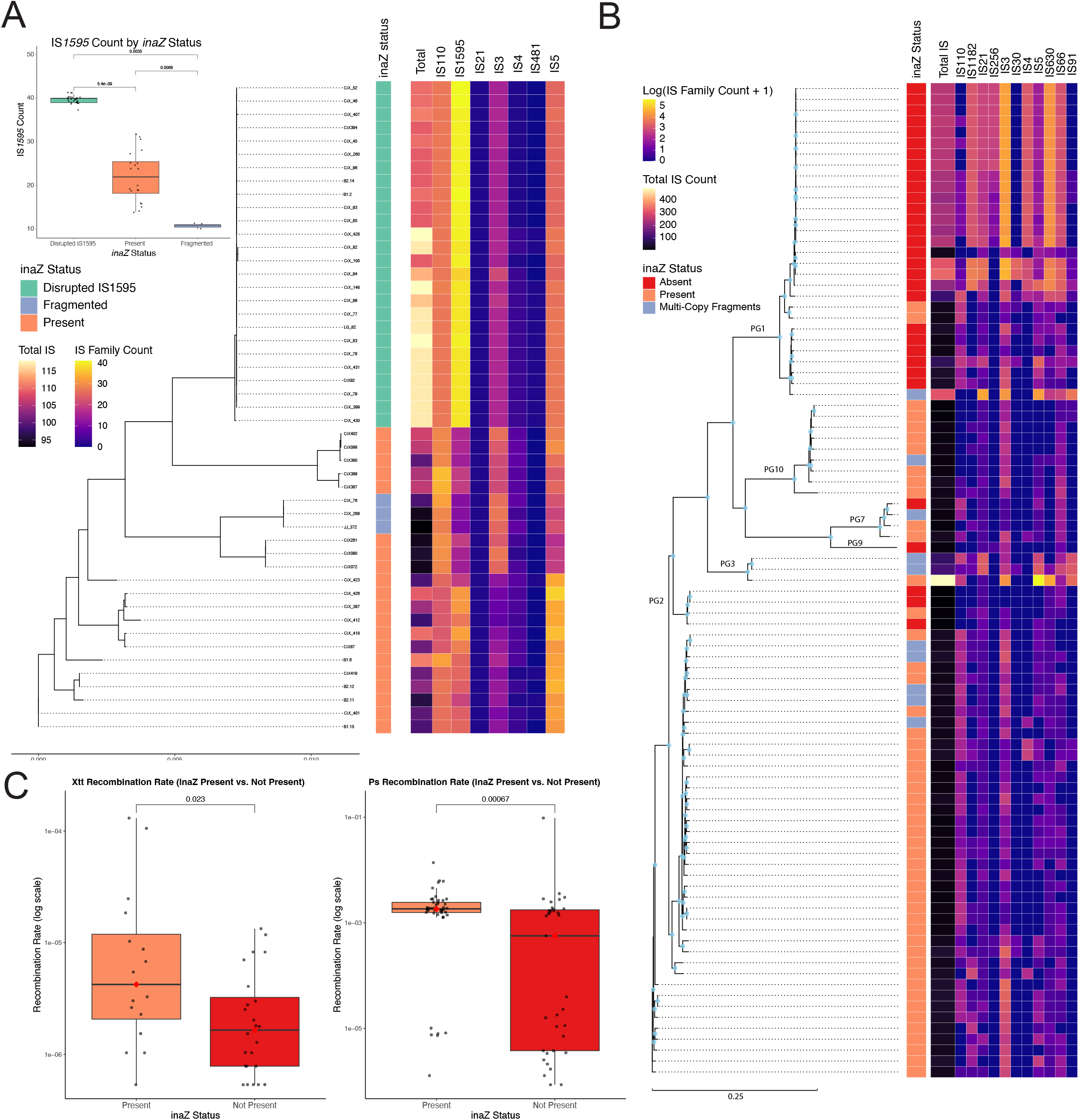
Lineages lacking potent InaZ are genetically isolated from the closely-related *inaZ*-containing bacteria. A&B) Maximum likelihood trees based on whole genome alignment represent the distribution of N. American Midwest *X. translucens* (A) and the metapopulation of *P. syringae* (B) that contain, lack or have mutated *inaZ. inaZ* status is denoted as present (orange), absent (red), fragmented (blue) or fragmented by IS1595 (green) as identified by ISEscan (52). C) Graphs quantify *X. translucens* and *P. syringae* recombination rates calculated with ClonalFrameML (64).

We created a novel *in silico* approach that predicts InaZ function (Fig. 1B). The *inaZ* allele has three functional domains: an N-terminus, central repeat domains (CRDs) and a C-terminus that comprise predictable repeat sequences, and each play fundamental roles for InaZ to be functional and are directly associated with the efficiency of INA (15–17). We observed that predicted non-functional InaZ homologs are strongly associated with either low amino acid alignment identity (e.g., < 60% identity) or the absence of essential functional components, defined as < 50% coverage of the N- or C-terminal domains or < 15 central repeat domains compared to reference InaZ proteins (UniProt accessions: P06620 for *Pseudomonas*, UPU47334.1 for *Xanthomonas*, and P20469 for *Pantoea*) (Fig. 1). These proteins with perturbed function are consistent with our work demonstrating that organisms lacking functional signatures in InaZ are reduced for INA under the tested conditions (no activity for 10e6 cells at -10°C) (18) (Fig. S3). Overall, these patterns suggest that this protein emerged in a Gammaproteobacteria common ancestor and that INA is an ancestral feature that is frequently lost as previously proposed (11, 19).

### INA limited bacteria are genetically isolated

For *P. syringae*, the link between INA and dispersal has been inferred from the observation that the frequency of INA strains in precipitation is significantly enriched compared to these frequencies in other substrates including clouds (8, 20). The importance of INA in global dispersal of this group of bacteria is further reinforced by the observation that the most ubiquitous genotypes of *P. syringae* are highly ice nucleation active (8). At present, these are the main observations that support the role of ice nucleation activity in facilitating global dispersal but the universality across Gammaproteobacteria remains to be confirmed. By assuming that loss of potent INA limits long-distance dispersal, here we tested the hypothesis that loss of INA in this group of bacteria leads to genetic isolation from related lineages.

We examined publicly available complete and chromosome level genomes for *P. syringae* (n=91) and sequenced genomes from a local population of *X. translucens* from the northern Midwest USA collected from a single plant species, *Hordeum vulgare* (n=48, 22 INA). This allowed for both generalizable conclusions on genome evolution across two species and in a respective local population with differing INA status from the same ecosystem.

The clades of *inaZ*-damaged or -lacking genomes were genetically similar appearing clonal and diverged from their closest relatives that contained *inaZ*. This suggested that these bacteria who would lack potent InaZ are genetically isolated (Fig. 3). These bacterial genomes, which contained versions of *inaZ* that were fragmented, disrupted by IS*1595* or completely absent, have ISE distributions distinct from *inaZ*-containing lineages (Fig. 3). As the expected corollary, we found that ice nucleation active bacteria had a significantly higher recombination rate than their ice nucleation inactive counterparts in both *X. translucens* (Wilcoxon, p = 0.023) and *Pseudomonas syringae* (Wilcoxon, p = 0.035) (Fig. 3). We observed a similar trend for the Brassica-associated lineage *X. campestris* pv. *campestris* (Xcc), in contrast to *X. campestris* as a whole. In Xcc, inaZ-lacking strains grouped separately from *inaZ*-containing lineages and exhibited a distinct expansion of ISEs (Fig. S6A & B). Furthermore, we determined that the only genomes (five of 155 total) that experienced increased recombination contained *inaZ* (Fig. S6C). This demonstrates that *inaZ*-containing strains are primary hotspots for genetic diversity in these populations.

To determine if the presence or absence of a functional *inaZ* gene was a random trait or a characteristic of specific evolutionary clonal lineages, we also tested for phylogenetic signal between genomes that lacked or contained *inaZ*. Notably in both species, the *inaZ* presence exhibited a remarkably strong correlation with the core genome phylogeny. For *X. translucens*, the phylogenetic signal was exceptionally strong (Pagel’s λ = 0.999; p < 0.001), and the trait was found to be highly conserved and grouped on the tree, significantly more than expected under a standard evolutionary model (Fritz & Purvis’ D = -0.56; p < 0.001). We observed a similarly strong and highly significant phylogenetic signal in *P. syringae* (Pagel’s λ = 0.987; p < 0.001), with the *inaZ* gene again showing a highly conserved, non-random distribution (Fritz & Purvis’ D = 0.11; p < 0.001).

### *inaZ* loss or mutation alters ecological behavior

As gene loss commonly accompanies niche differentiation in plant-associated bacteria (*22, 23*), our research demonstrates, to our knowledge, a keystone example where the loss of a biophysical process drives transmission changes in microorganisms. The Gammaproteobacteria examined here are plant-associated or pathogenic. As such, they can exploit a range of processes for dispersal in association with plant tissues and via aerial dispersal and movement with surface waters depending on their capacity to survive (8, 14, 23–25) (Fig. S7). In light of these observations, we posit that bacteria that lost *inaZ* have lost or inefficient INA and a marked dependence on transmission by plant tissues or insects and a notably increased ecological signature of isolation.

For example, lineages that lost *inaZ* include *X. translucens, X. campestris, Xanthomonas vasicola, Xanthomonas oryzae, P. syringae* pv. actidineae, which are known to be dispersed by plant seed or propagated cuttings, and *Pantoea stewartii*, which is vectored by beetles (Fig. 2) (23). The global movement of these organisms has instead resulted from well-documented human-driven transmission with plant material or insects as a key epidemiological process (26– 29). *P. syringae* pv. actidineae, *X. vasicola* and *X. oryzae* lack *inaZ* and have been recently moved across continents where they were not previously present by human movement of cuttings or seed (26–30).This does not imply that INA positive bacteria do not have access to these transmission processes; as illustrated in Fig. S7, we highlight that mixing by long-distance atmospheric dispersal confounds other forms of transmission (e.g. plant tissues).

The non-uniform distribution of *inaZ* loss events suggests that within species, *inaZ-*containing populations may persist (Fig. 1&2), but loss of *inaZ* may lead to localized, region-specific and endemic populations (Fig. 3C). Atmosphere-dispersed populations may provide a reservoir for continual introduction to new environments, while clonally distributed, *inaZ-*lacking populations may face extinction in their restricted environments or fail to propagate into new environments at the rate of *inaZ-*containing relatives.

### Concluding remarks

This work raises questions about the range of consequences of *inaZ* loss. Here we demonstrated that loss of INA or reduction of its potency alters microbial evolution, genetic diversity and dispersal. These ultimately have potential impacts on crop health and its management in spite of the evolutionary history of INA being independent from crop production (19). The second potential consequence is on Earth system processes. The warm temperature activity of bacterial ice nuceli give them a potentially greater role in the water cycle in a changing climate where the changing temperature profiles of clouds could reduce the opportunities for the abundant mineral ice nuclei to set off rainfall (31). A key challenge will be to discover if there are anthropogenic drivers of INA loss in bacteria and to determine their relative importance compared to natural drivers. A critical step in identifying drivers of loss will be in dating the time of emergence of lines of bacteria that have lost *inaZ*.

### Materials and Methods

#### Phylogenomic and phylogenetic analysis across *inaZ*-containing bacteria

Metadata for all genome assemblies at NCBI from the genera *Xanthomonas, Pseudomonas*, and *Pantoea* was downloaded on 2023-10-17 with the “summary genome taxon” command from the NCBI datasets tool v. 15.16.1 (Table S1 & Data S1) (32). Among the 36,430 available genome assemblies, we used the downloaded metadata to select those that fulfilled the following criteria: labelled as “Complete Genome” by NCBI, consisting of at most 3 contigs, having taxonomy assigned to at least the species level, and having been sequenced with technology that included Illumina and/or Pacific Biosciences. The resulting selection of 1,531 assemblies was downloaded with the NCBI datasets “download genome accession” command on 2023-10-28. The downloaded assemblies were annotated with Prokka v. 1.14.6 (33), and a custom BLAST database was created with all resulting proteomes (“.faa” FASTA files) using the “makeblastdb” command from the BLAST+ suite v. 2.13.0 (34).

A focal InaZ protein sequence of 1,567 amino acids from *Xanthomonas translucens* pv. *undulosa* isolate UPB513 (GCF_023221635.1) was downloaded from NCBI with the command “efetch -db protein -format fasta -id UPU47334.1” from the NCBI Entrez Direct toolkit v. 16.2 (35). A BLAST search for this protein sequence against the abovementioned custom BLAST database was performed using the “blastp” tool from the BLAST+ suite. The resulting hits were filtered by e-value (<1×10^-6^) and by percent identity (>29%), and parsed to produce a first-pass list of 231 genomes with one or more potential copies of the InaZ gene.

Orthofinder v. 2.5.5 (36) was run using “-M msa” and otherwise with default options to infer orthologs among the selected 231 genomes with one or more potential InaZ gene copies and to produce a species tree using sequences from all orthogroups which had all included genomes represented in them. Separate nucleotide and protein gene trees were created for the orthogroup containing the focal InaZ protein from UPB513 using IQ-tree v. 2.2.2.7 (37, 38). The protein tree was created directly from the alignment produced by Orthofinder, whereas the corresponding nucleotide alignment was produced with MAFFT v. 7.520 (39) after extracting the nucleotide sequence for each gene in the focal orthogroup using the genomic coordinates for each gene in corresponding genome’s GFF file with the “getfasta” command from Bedtools v. 2.31.0 (40). The resulting trees were plotted using ggtree v. 3.8.2 (41) in R v. 4.3.0 (42).

#### InaZ protein subdomain analysis

To discriminate between functional and non-functional *inaZ* loci, we extracted the N-terminus, C-terminus, and central repeat domains (CRDs) associated with each *inaZ* hit. We identified these subdomain sequences by extracting the CRD and then circumscribing the up- and downstream sequences to their respective subdomain terminus. We identifed CRD sequences by identifying an anchor CRD amino acid sequence that hits the reported conserved CRD sequence: “GYGST*TA***S*L[T/I]A”. We then extended the overall CRD by identifying surrounding CRD sequences with a maximum hamming distance of 4 relative the conserved CRD query. With CRDs in hand, the N- and C-terminus were extracted from the surrounding region, and the percent identity of each terminus was obtained by *BLASTp* against a previously published *Ina* terminus for the respective genus: *Pantoea InaA* Panan_P20469 (UniProt), *Pseudomonas* Psesy_P06620 (UniProt), and *Xanthomonas* Xantra_UPU47334.1 (GenBank).

#### Single-copy ortholog phylogenomic tree for *inaZ* synteny analysis

To infer the evolution of the *inaZ* gene among the genomes that contain a homolog, we built a phylogenomic tree derived from well-supported single-copy orthologs. The initial dataset of 1,530 genomes was intractable for use with OrthoFinder, so to determine single-copy orthologs (SCOs) we extracted up to two representative genomes of each species in the initial dataset and submitted this truncated dataset to OrthoFinder (36). This method identified 171 total SCOs. We then retrieved orthologs of these SCOs across the entire dataset by building hidden-Markov models (HMMs) of the *mafft –*derived multiple sequence alignments of each SCO and retrieving the best hit of each HMM in the entire dataset (43). To determine which SCOs are informative for phylogenomic analysis, we extracted 52 (30.4%) of SCOs that had individual gene trees with greater than 25% average support. Individual SCO trees were obtained by aligning each group of SCO sequences using *mafft*, trimming sequences via *ClipKIT*, determining the best-fit evolutionary model using *ModelFinder*, and reconstructing phylogenies using *IQ-TREE* (37, 38, 44–46). The final phylogenomic tree was built from the 52 well-supported SCO alignments using a multi-gene partition model that individually applies the best-fit evolutionary model previously determined for the SCO to the partition of the concatenated alignment that corresponds to that set of sequences using IQ-TREE (37, 38).

#### inaZ synteny analysis

To compare synteny among *inaZ* loci and infer the causal agents of InaZ loss, we extracted loci that contain functional *inaZ* genes and identified homologous loci that lack inaZ or have a putative pseudogene of *inaZ*. To adequately sample allelic diversity while maintaining a visualizable synteny diagram, we randomly selected a representative genome from each clade of functional InaZ, putative non-functional InaZ, and InaZ-absent loci referencing the single-copy ortholog phylogenomic tree. We represented functional inaZ loci by extracting 5 kB up- and downstream of functional *inaZ* genes. We identified homologous loci in *inaZ*-absent genomes by querying a closely related functional locus using the Cluster Reconstruction and Phylogenetic Analysis Pipeline (CRAP) as implemented in mycotools(47, 48). This method is implemented by querying each gene in a query locus via *BLASTp*, extracting 5 kB up- and downstream of hits, truncating the resulting hit locus boundaries to the furthest genes that share homology to the query locus, and identifying the most similar locus for each hit genome. Similarity between hit and query locus is quantified by the Jaccard index of the presence-absence of homologous genes between the query and subject scaled by the average percent identity of overlapping genes (48). We then visualized the shared synteny between extracted homologous loci using *clinker* (49).

#### X. translucens ISE profiling

High-quality, complete and contiguous assemblies for 39 strains of *X. translucens* belonging to 12 pathovars, with one strain not identified to the pathovar level were independently annotated with Prokka v1.14.6 and 167 orthologous genes shared among all genomes were identified and aligned with Roary v3.13.0 (cite Prokka and Roary) (33, 50). A Bayesian phylogenetic tree was generated from this alignment using MrBayes v3.2.7a (cite MrBayes) (51) and the tree was visualized with FigTree v1.4.4 (available at https://tree.bio.ed.ac.uk/software/figtree/). ISEScan v1.7.2.3 was used to identify, classify and count ISEs in each genome (52). The numbers of ISEs belonging to families IS110, IS1595, IS3, IS4, IS5, and Other were plotted as circles with the plotrix R package (53) with ISE count dictating circle radii. Total ISE counts were plotted in R with the barplot function.

#### Intraspecies recombination analysis

Genome assemblies for *Pseudomonas syringae, Xanthomonas translucens*, and *Xanthomonas campestris pv. campestris*were retrieved from the National Center for Biotechnology Information (NCBI) database. The NCBI Datasets command-line tool (v13.2.0) was utilized to perform the downloads. For each species, a targeted query was used to select genomes with an assembly level of “complete” or “chromosome” to ensure high-quality, closed reference genomes were used for the analysis. All downloaded genome assemblies were subsequently re-annotated using a uniform method to avoid biases from different annotation pipelines. Genome annotation was performed with Prokka (v1.14.6), using the --kingdom Bacteriaflag and default parameters. This process generated standardized GenBank-format (.gff) files for each genome, which served as the input for all downstream pangenome and comparative analyses.

For each species, a pangenome analysis was conducted using Roary (v3.13.0) on the complete set of Prokka-generated .gff files. Roary was run with the -e --mafft options to create a core gene alignment from the single-copy core genes present in at least 99% of the isolates. The resulting core gene alignment file (core_gene_alignment.aln) was used to construct a maximum-likelihood phylogenetic tree for each species. Tree inference was performed with IQ-TREE (v2.1.2). The optimal substitution model was determined for each alignment using ModelFinder Plus (-m MFP), and branch support was assessed using 1000 ultrafast bootstrap replicates (-B 1000). The final, unrooted tree with the highest likelihood was taken from the .treefile output for downstream analyses.

To quantify recombination rates and events within each species, we used ClonalFrameML (v1.12)(54). For each species, the core genome alignment from Roary and the corresponding unrooted phylogenetic tree from IQ-TREE were used as input. ClonalFrameML was run with default parameters to infer a recombination-aware phylogeny and calculate key recombination parameters. The recombination rate for each strain (or “tip” in the tree) was extracted from the posterior mean of the R/thetaparameter in the .em.txt output file. Strains for which ClonalFrameML did not infer a rate were assigned a recombination rate of zero, representing no detectable recombination. The total number of recombination events per strain was calculated by counting the occurrences of each strain’s ID in the “Node” column of the importation_status.txt output file. ISEs were identified and classified in each genome assembly using ISEScan (v1.7.2.3) with default parameters (55). The .tsv output files from each individual ISEScan run were parsed and aggregated using a custom Python script utilizing the pandas library(55). This script generated a summary matrix for each species, detailing the total count of each identified IS family per genome, as well as the total IS element abundance for each strain. All statistical analyses and data visualizations were performed in R (v4.2.1; R Core Team 2021). Data manipulation and aggregation were conducted using the dplyr (https://dplyr.tidyverse.org) and tidyr (https://tidyr.tidyverse.org) packages. All figures were generated using ggplot2 (https://ggplot2.tidyverse.org) and the ggtree (https://guangchuangyu.github.io/software/ggtree/) package for phylogenetic visualizations. Multi-panel figures were assembled using the patchwork library. Comparisons of recombination rates and IS element counts between inaZ+ and *inaZ*-groups were performed using a non-parametric Mann-Whitney U test (Wilcoxon rank-sum test), implemented with the ggpubr (https://rpkgs.datanovia.com/ggpubr/) package. For cases where one group exhibited zero variance (e.g., no recombination in any *inaZ*-*Xcc* strains), a Fisher’s Exact Test was used to test for a significant association between *inaZ* status and the presence or absence of recombination. To test if the *inaZ* gene presence was significantly associated with clonal lineages, we tested for phylogenetic signal using the phytools and caper packages (56). We calculated two metrics: Pagel’s Lambda (λ) and Fritz & Purvis’s D-statistic (57–60).

#### Rain microbiome analysis

Rainwater was collected over 3 rain events in June 2025 in 10and transferred into large bags. A 2 L of liquid was filtered through a 0.45 µm sterile membrane. The filters were sonicated in 10 mL of CTAB buffer (composition per liter: 8 g CTAB, 10 g SDS, 25 g mannitol, 47 g NaCl, 2 g Na_2_EDTA, 10 g PVPP) and incubated for 60 minutes at 65 °C. One milliliter of the CTAB solution was then transferred to 2 mL tubes, followed by the addition of 1 mL chloroform. Samples were vortexed for 5 minutes and centrifuged at maximum speed for 10 minutes. The upper aqueous phase was recovered, and two volumes of ethanol were added. Tubes were stored at –20 °C overnight, then centrifuged at maximum speed for 10 minutes at 4 °C. DNA pellets were resuspended in water and incubated at 50 °C for 1 hour to ensure complete dissolution. A total of 20 ng of DNA was sequenced at the Applied Microbiology Services Laboratory (AMSL), The Ohio State University using the Illumina DNA prep library and the NextSeq2000. Raw reads were trimmed using Trimmomatic v0.39. The cleaned reads from Trimmomatic were taxonomically classified using Kaiju (61), employing a custom database. This database includes all complete *inaZ* protein sequences from Gammaproteobacteria, downloaded from UniProt. Classification results were visualized using krona (62).

#### Ice nucleation activity

We assessed ice nucleation activity using two approaches. We used the immersion freezing assay on droplets of aqueous bacterial suspensions as described (8). In this assay, droplets containing a standardized total of 10^7^ CFU were tested for freezing events at temperatures at – 6°C. Ice nucleation experiments were also performed using the high-throughput Twin-plate Ice Nucleation Assay (TINA), which has been described in detail elsewhere (63). In a typical experiment, the investigated IN sample was serially diluted 10-fold by a liquid handling station (epMotion ep5073, Eppendorf, Hamburg, Germany). 96 droplets (droplet volume: 3 μL) per dilution were placed on two 384-well plates and tested with a continuous cooling-rate of 1 °C/min from 0 to −30 °C. The droplet freezing events were detected by two infrared cameras (Seek Thermal Compact XR, Seek Thermal Inc., Santa Barbara, CA). The uncertainty in the temperature of the setup was ± 0.2 °C (63).

## Supporting information

BLASTData

## Acknowledgments

We would like to acknowledge access to high performance computing resources provided by the Ohio Supercomputer Center. Google Gemini was used for article editing and creating code. The authors are grateful for funding supported by: The Ohio State University’s Infectious Diseases Institute (JMJ and VRR); USDA NIFA through the NSF/NIFA Plant Biotic Interactions Program award no. 2018-67013-28490 (JMJ); US Fulbright Scholar Award to Uruguay (JMJ); USDA NIFA Postdoctoral Fellowship award no. 2018-08122 (SPC); IMPLANTEUS (JMJ, CM).

## Notes

### Competing Interest Statement

The authors have declared no competing interest.

## References

1. S. E. Lindow, D. C. Arny, C. D. Upper, Bacterial Ice Nucleation: A Factor in Frost Injury to Plants 1. Plant Physiology 70, 1084–1089 (1982).

2. B. M. Wiegmann, et al., Episodic radiations in the fly tree of life. Proceedings of the National Academy of Sciences 108, 5690–5695 (2011).

3. G. Ruaux, S. Lumineau, E. de Margerie, The development of flight behaviours in birds. Proceedings of the Royal Society B: Biological Sciences 287, 20200668 (2020).

4. L. Yafetto, et al., The Fastest Flights in Nature: High-Speed Spore Discharge Mechanisms among Fungi. PLOS ONE 3, e3237 (2008).

5. N. A. Wright, D. W. Steadman, C. C. Witt, Predictable evolution toward flightlessness in volant island birds. Proceedings of the National Academy of Sciences 113, 4765–4770 (2016).

6. V. E. Kutschera, et al., Purifying Selection in Corvids Is Less Efficient on Islands. Mol Biol Evol 37, 469–474 (2020).

7. C. E. Morris, Leyronas, C., Nicot, P.C., “Movement of Bioaerosols in the Atmosphere and the Consequences for Climate and Microbial Evolution (Chapter 16)” in Aerosol Science: Technology and Applications, Colbeck and Lazaridis, (John Wiley & Sons, Ltd), pp. 393–416.

8. C. E. Morris, et al., The life history of the plant pathogen Pseudomonas syringae is linked to the water cycle. The ISME Journal (2008). 10.1038/ismej.2007.113.

9. S. Lindow, History of Discovery and Environmental Role of Ice Nucleating Bacteria. Phytopathology® 113, 605–615 (2023).

10. B. C. Christner, C. E. Morris, C. M. Foreman, R. Cai, D. C. Sands, Ubiquity of Biological Ice Nucleators in Snowfall. Science 319, 1214–1214 (2008).

11. P. K. Wolber, “Bacterial Ice Nucleation” in Advances in Microbial Physiology, A. H. Rose, Ed. (Academic Press, 1993), pp. 203–237.

12. S. E. Lindow, J. H. J. Leveau, Phyllosphere microbiology. Current Opinion in Biotechnology 13, 238–243 (2002).

13. C. E. Morris, D. G. Georgakopoulos, D. C. Sands, Ice nucleation active bacteria and their potential role in precipitation. J. Phys. IV France 121, 87–103 (2004).

14. M. E. Mechan Llontop, et al., Exploring Rain as Source of Biological Control Agents for Fire Blight on Apple. Front. Microbiol. 11 (2020).

15. S. Hartmann, et al., Structure and Protein-Protein Interactions of Ice Nucleation Proteins Drive Their Activity. Frontiers in Microbiology Volume 13–2022 (2022).

16. G. Renzer, et al., Hierarchical assembly and environmental enhancement of bacterial ice nucleators. Proceedings of the National Academy of Sciences 121, e2409283121 (2024).

17. J. Forbes, et al., Water-organizing motif continuity is critical for potent ice nucleation protein activity. Nat Commun 13, 5019 (2022).

18. C. E. Morris, et al., Inferring the evolutionary history of the plant pathogen Pseudomonas syringae from its biogeography in headwaters of rivers in North America, Europe and New Zealand. mBio (2010). 10.1128/mBio.00107-10.Editor.

19. C. E. Morris, et al., Bioprecipitation: a feedback cycle linking Earth history, ecosystem dynamics and land use through biological ice nucleators in the atmosphere. Global Change Biology 20, 341–351 (2014).

20. M. Joly, et al., Ice nucleation activity of bacteria isolated from cloud water. Atmospheric Environment 70, 392–400 (2013).

21. E. Gluck-Thaler, et al., Repeated gain and loss of a single gene modulates the evolution of vascular plant pathogen lifestyles. Science Advances 6, eabc4516 (2020).

22. R. A. Melnyk, S. S. Hossain, C. H. Haney, Convergent gain and loss of genomic islands drive lifestyle changes in plant-associated Pseudomonas. The ISME Journal 13, 1575–1588 (2019).

23. M.-A. Jacques, et al., Using Ecology, Physiology, and Genomics to Understand Host Specificity in Xanthomonas. Annu. Rev. Phytopathol. 54, 163–187 (2016).

24. B. Dutta, et al., Transmission of Pantoea ananatis and P. agglomerans, Causal Agents of Center Rot of Onion (Allium cepa), by Onion Thrips (Thrips tabaci) Through Feces. 10.1094/PHYTO-07-13-0199-R (2014). Available at: https://apsjournals.apsnet.org/doi/10.1094/PHYTO-07-13-0199-R [Accessed 30 September 2025].

25. C. E. Morris, J. R. Lamichhane, I. Nikolić, S. Stanković, B. Moury, The overlapping continuum of host range among strains in the Pseudomonas syringae complex. Phytopathology Research 1, 4 (2019).

26. I. Donati, et al., Pseudomonas syringae pv. actinidiae: Ecology, Infection Dynamics and Disease Epidemiology. Microbial Ecology 80, 81–102 (2020).

27. S. L. Arias, et al., Occurrence in Seeds and Potential Seed Transmission of Xanthomonas vasicola pv. vasculorum in Maize in the United States. Phytopathology® 110, 1139–1146 (2020).

28. A. L. Perez-Quintero, et al., Genomic Acquisitions in Emerging Populations of Xanthomonas vasicola pv. vasculorum Infecting Corn in the United States and Argentina. Phytopathology® 110, 1161–1173 (2020).

29. V. Schepler-Luu, et al., Genome editing of an African elite rice variety confers resistance against endemic and emerging Xanthomonas oryzae pv. oryzae strains. eLife 12, e84864 (2023).

30. H. C. McCann, et al., Origin and Evolution of the Kiwifruit Canker Pandemic. Genome Biology and Evolution 9, 932–944 (2017).

31. H. Luo, et al., Robust variation trends in cloud vertical structure observed from three-decade radiosonde record at Lindenberg, Germany. Atmospheric Research 281, 106469 (2023).

32. N. A. O’Leary, et al., Exploring and retrieving sequence and metadata for species across the tree of life with NCBI Datasets. Sci Data 11, 732 (2024).

33. T. Seemann, Prokka: rapid prokaryotic genome annotation. Bioinformatics 30, 2068–2069 (2014).

34. C. Camacho, et al., BLAST+: architecture and applications. BMC Bioinformatics 10, 421 (2009).

35. J. Kans, “Entrez direct: E-utilities on the UNIX command line” in Entrez Programming Utilities Help [Internet], (National Center for Biotechnology Information (US), 2024).

36. D. M. Emms, S. Kelly, OrthoFinder: phylogenetic orthology inference for comparative genomics. Genome Biology 20, 238 (2019).

37. B. Q. Minh, et al., IQ-TREE 2: New Models and Efficient Methods for Phylogenetic Inference in the Genomic Era. Molecular Biology and Evolution 37, 1530–1534 (2020).

38. L.-T. Nguyen, H. A. Schmidt, A. von Haeseler, B. Q. Minh, IQ-TREE: A Fast and Effective Stochastic Algorithm for Estimating Maximum-Likelihood Phylogenies. Mol Biol Evol 32, 268–274 (2015).

39. K. Katoh, K. Misawa, K. Kuma, T. Miyata, MAFFT: a novel method for rapid multiple sequence alignment based on fast Fourier transform. Nucleic Acids Research 30, 3059–3066 (2002).

40. A. R. Quinlan, I. M. Hall, BEDTools: a flexible suite of utilities for comparing genomic features. Bioinformatics 26, 841–842 (2010).

41. G. Yu, D. K. Smith, H. Zhu, Y. Guan, T. T.-Y. Lam, ggtree: an r package for visualization and annotation of phylogenetic trees with their covariates and other associated data. Methods in Ecology and Evolution 8, 28–36 (2017).

42. R Core Team, R: A Language and Environment for Statistical Computing (R Foundation for Statistical Computing, 2024).

43. M. Madera, J. Gough, A comparison of profile hidden Markov model procedures for remote homology detection. Nucleic Acids Res 30, 4321–4328 (2002).

44. J. L. Steenwyk, T. J. B. Iii, Y. Li, X.-X. Shen, A. Rokas, ClipKIT: A multiple sequence alignment trimming software for accurate phylogenomic inference. PLOS Biology 18, e3001007 (2020).

45. K. Katoh, K. Misawa, K. Kuma, T. Miyata, MAFFT: a novel method for rapid multiple sequence alignment based on fast Fourier transform. Nucleic Acids Res 30, 3059–3066 (2002).

46. S. Kalyaanamoorthy, B. Q. Minh, T. K. F. Wong, A. von Haeseler, L. S. Jermiin, ModelFinder: fast model selection for accurate phylogenetic estimates. Nat Methods 14, 587–589 (2017).

47. Z. Konkel, J. C. Slot, Mycotools: An Automated and Scalable Platform for Comparative Genomics. Preprint] (2023). Available at: https://www.biorxiv.org/content/10.1101/2023.09.08.556886v1 [Accessed 13 November 2025].

48. Z. Konkel, L. Kubatko, J. C. Slot, CLOCI: unveiling cryptic fungal gene clusters with generalized detection. Nucleic Acids Res 52, e75 (2024).

49. C. L. M. Gilchrist, Y.-H. Chooi, clinker & clustermap.js: automatic generation of gene cluster comparison figures. Bioinformatics 37, 2473–2475 (2021).

50. A. J. Page, et al., Roary: rapid large-scale prokaryote pan genome analysis. Bioinformatics 31, 3691–3693 (2015).

51. F. Ronquist, et al., MrBayes 3.2: Efficient Bayesian Phylogenetic Inference and Model Choice Across a Large Model Space. Systematic Biology 61, 539–542 (2012).

52. Z. Xie, H. Tang, ISEScan: automated identification of insertion sequence elements in prokaryotic genomes. Bioinformatics 33, 3340–3347 (2017).

53. J. Lemon, Plotrix: A Package in the Red Light District of R. R-News 8–12 (2006).

54. X. Didelot, D. J. Wilson, ClonalFrameML: Efficient Inference of Recombination in Whole Bacterial Genomes. PLOS Computational Biology 11, e1004041 (2015).

55. Z. Xie, H. Tang, ISEScan: automated identification of insertion sequence elements in prokaryotic genomes. Bioinformatics 33, 3340–3347 (2017).

56. L. J. Revell, phytools 2.0: an updated R ecosystem for phylogenetic comparative methods (and other things). PeerJ 12, e16505 (2024).

57. S. A. Fritz, A. Purvis, Selectivity in Mammalian Extinction Risk and Threat Types: a New Measure of Phylogenetic Signal Strength in Binary Traits. Conservation Biology 24, 1042–1051 (2010).

58. D. Barker, M. Pagel, Predicting Functional Gene Links from Phylogenetic-Statistical Analyses of Whole Genomes. PLoS Computational Biology 1, e3–e3 (2005).

59. W. D. Pearse, T. J. Davies, E. M. Wolkovich, How to Define, Use, and Interpret Pagel’s λ (Lambda) in Ecology and Evolution. Global Ecology and Biogeography 34, e70012 (2025).

60. M. Pagel, A. Meade, D. Barker, Bayesian estimation of ancestral character states on phylogenies. Systematic Biology (2004). 10.1080/10635150490522232.

61. P. Menzel, K. L. Ng, A. Krogh, Fast and sensitive taxonomic classification for metagenomics with Kaiju. Nature Communications 7, 11257 (2016).

62. B. D. Ondov, N. H. Bergman, A. M. Phillippy, Interactive metagenomic visualization in a Web browser. BMC Bioinformatics 12, 385 (2011).

63. A. T. Kunert, et al., Twin-plate Ice Nucleation Assay (TINA) with infrared detection for highthroughput droplet freezing experiments with biological ice nuclei in laboratory and field samples. Atmospheric Measurement Techniques 11, 6327–6337 (2018).

64. X. Didelot, D. J. Wilson, ClonalFrameML: Efficient Inference of Recombination in Whole Bacterial Genomes. PLOS Computational Biology 11, e1004041 (2015).

