## SupplementalInformation for "Loss of dispersal via ice nucleation activity constrains microbial evolution"

**This PDF file includes:**

Supporting text

Figures S1 to S7

Tables S1 to S2

Legends for Datasets S1

SI References

**Other supporting materials for this manuscript include the following:**

Table S1 to S2

Datasets S1

Supporting Information Text

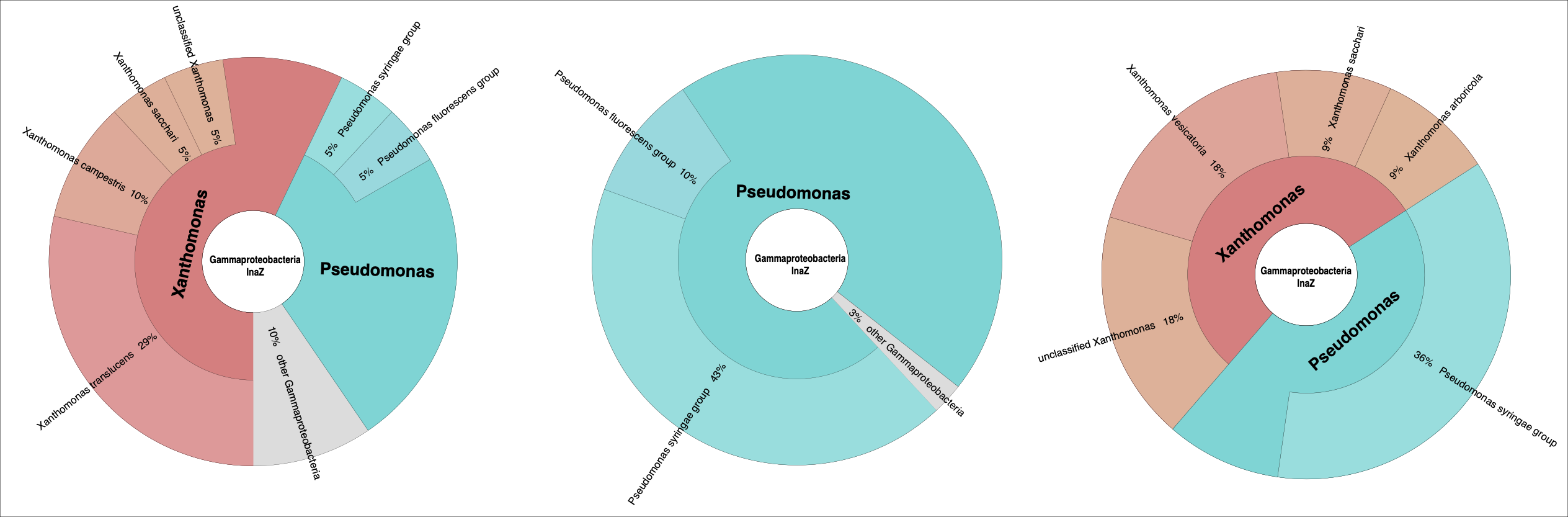

Fig. S1. InaZ is a protein function in rain metagenomes. Metagenome sequencing was carried out with NextSeq2000 from rain collected from in Columbus, Ohio. Uniprot database was used as a reference for mapping reads to a consensus inaZ sequence from Xanthomonas and Pseudomonas spp. Xanthomonas spp. and Pseudomonas spp. were identified in red/orange and turquoise, respectively.

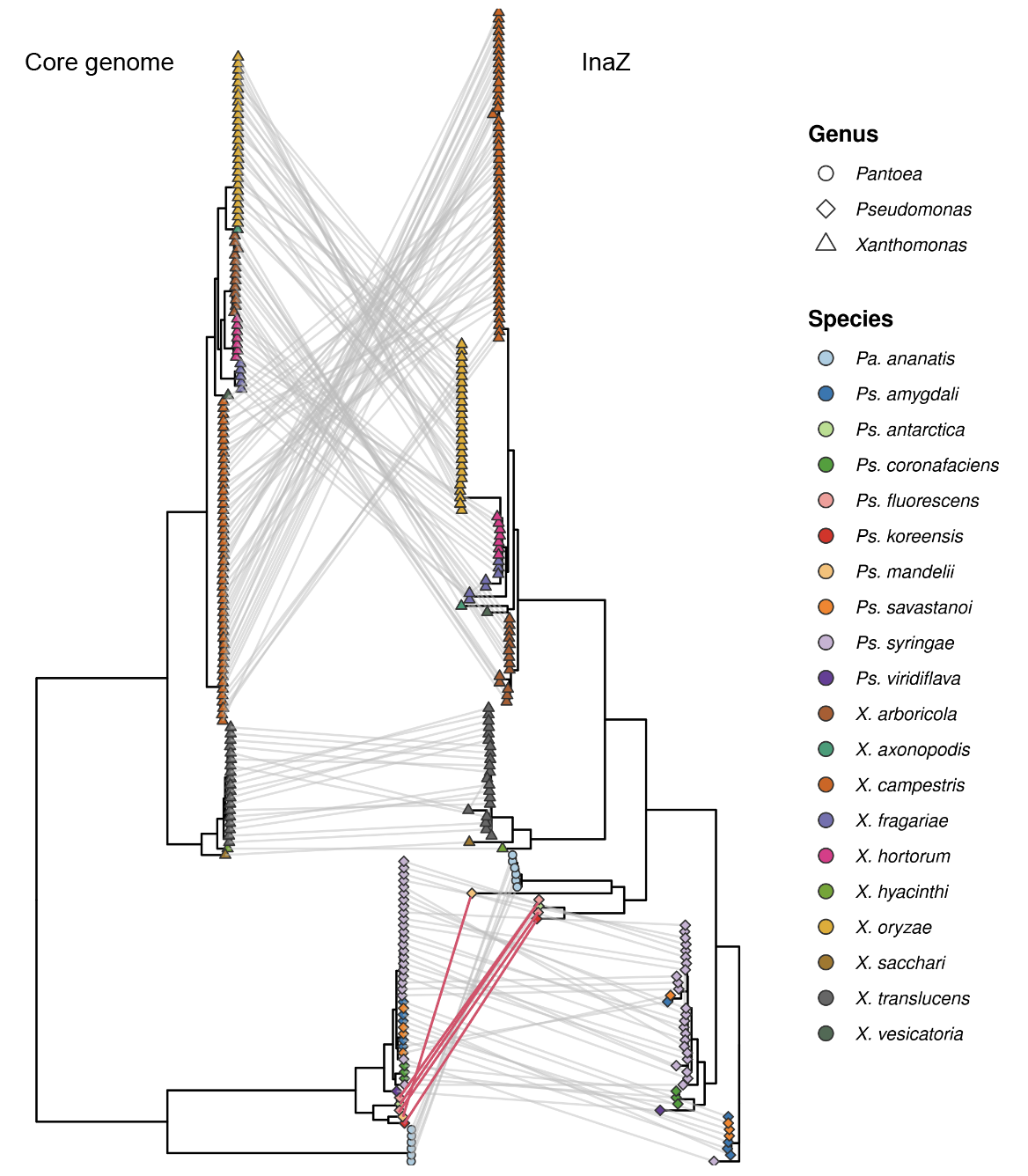

Fig. S2. InaZ protein tree parallels core genome phylogeny with potential horizontal transfer in *Xanthomonas*, *Pseudomonas* and *Pantoea*. Red lines indicate incongruencies where *inaZ* was hypothesized to be horizontally transferred.

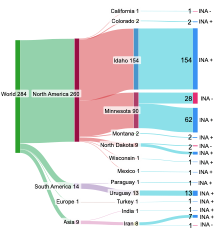

Fig. S3. INA from 284 Isolates of barley *X. translucens* isolates via immersion freezing (1). This sandkey diagram displays the INA of isolates collected across four continents. INA+ describes INA at -6^o^C; while INA- describes bacteria that did not freeze at -6^o^C.

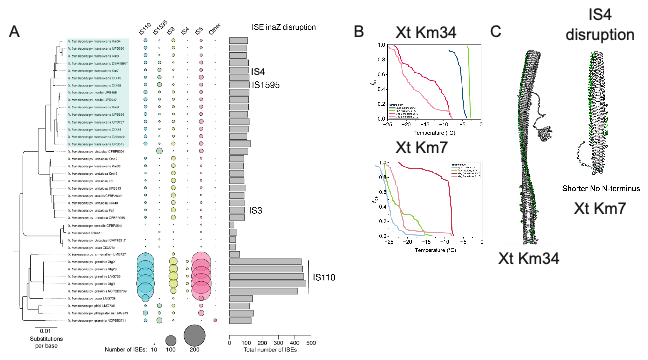

Fig. S4. ISEs have shaped the evolution of *X. translucens.* Phylogenetic tree of 39 strains of *X. translucens* inferred from alignment of 167 core genes using Bayesian inference (posterior probability = 1.00 for all branches, distance scale = substitutions per base). Number of ISEs per family (IS110, IS1595, IS3, IS4, IS5, and all other families) and total number of ISEs per strain are indicated by circle radii and barplots, respectively. ISE-mediated disruption of *inaZ* family indicated (e.g. IS1595 disruption of *inaZ* in CIX95, etc.). B) Ice nucleation activity was tested via TINA for strains Km33 and Km7. C) Alphafold3 prediction modeling of a wildtype protein amd mutated variant from strains Km33 and Km7. B&C) Km33 is a wild-type protein and enables freezing at -2C; Km7 was disrupted by IS4 removing the N-terminus and shortening the CRD domain and thus reducing INA.

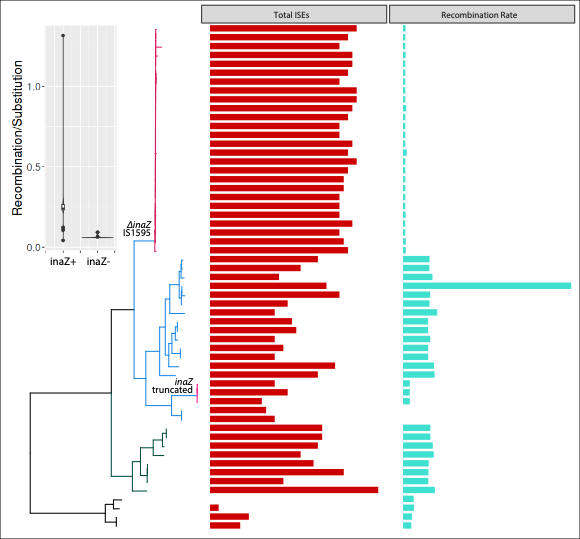

Fig. S5. Ice nucleation inactive lineages of X. translucens are genetically isolated from the global population. A) Single copy ortholog maximum likelihood phylogenomic trees represent the diversity of complete genome sequences of inaZ positive and negative (dark red: IS1595 disruption, light red: frameshift mutation). Bar graphs represent either ISE abundance (red) and recombination rate (turquoise).

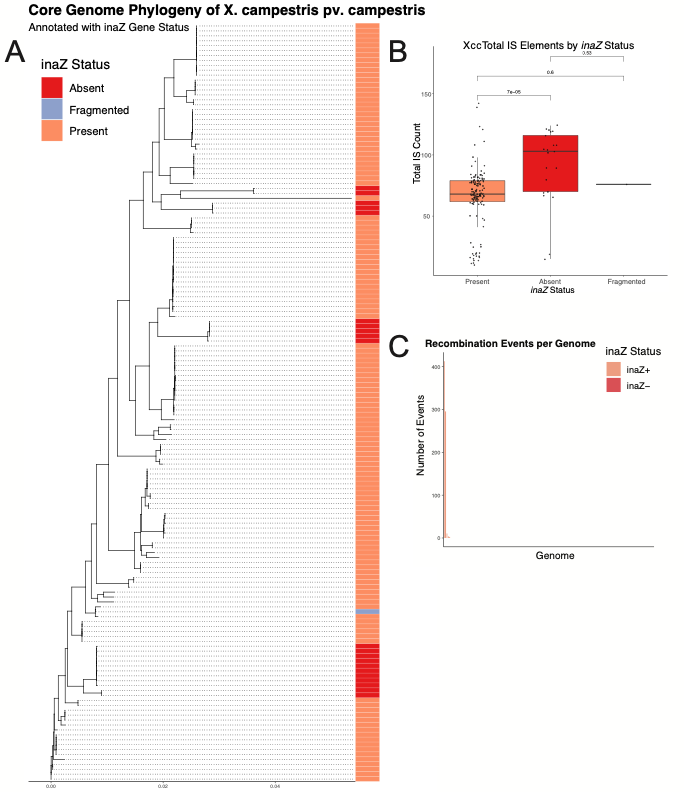

Fig. S6. Loss of *inaZ* in *Xanthomonas campestris* pv. campestris (Xcc) is linked to distinct phylogenetic lineages with higher insertion sequence abundance and reduced recombination. A) A maximum-likelihood core genome phylogeny was constructed from a concatenated alignment of core genes identified by Roary, with the tree inferred using IQ-TREE. Aligned to the tips of the tree is a heatmap indicating the status of the inaZ gene for each strain. Strains with an intact *inaZ* gene (status 1) are shown in orange, those where the gene is absent (status 0) are in red, and the strain with a fragmented gene (status 2) is highlighted in light blue. The tree demonstrates that the loss of *inaZ* is not random but is characteristic of specific clonal lineages. B) Boxplot comparing the distribution of the total number of insertion sequence elements (ISEs), quantified using ISEScan, across the three *inaZ* status groups. The plot shows a notable expansion in the number of ISEs in lineages where *inaZ* is absent (red) or fragmented (light blue) compared to lineages where it is present (orange). (C) Boxplot comparing the number of recombination events per genome, as inferred by ClonalFrameML from the core genome alignment. Strains are grouped by their *inaZ* phenotype (*inaZ*-positive vs. *inaZ*-negative, which combines absent and fragmented statuses). The comparison highlights differences in the extent of recombination between the two groups.

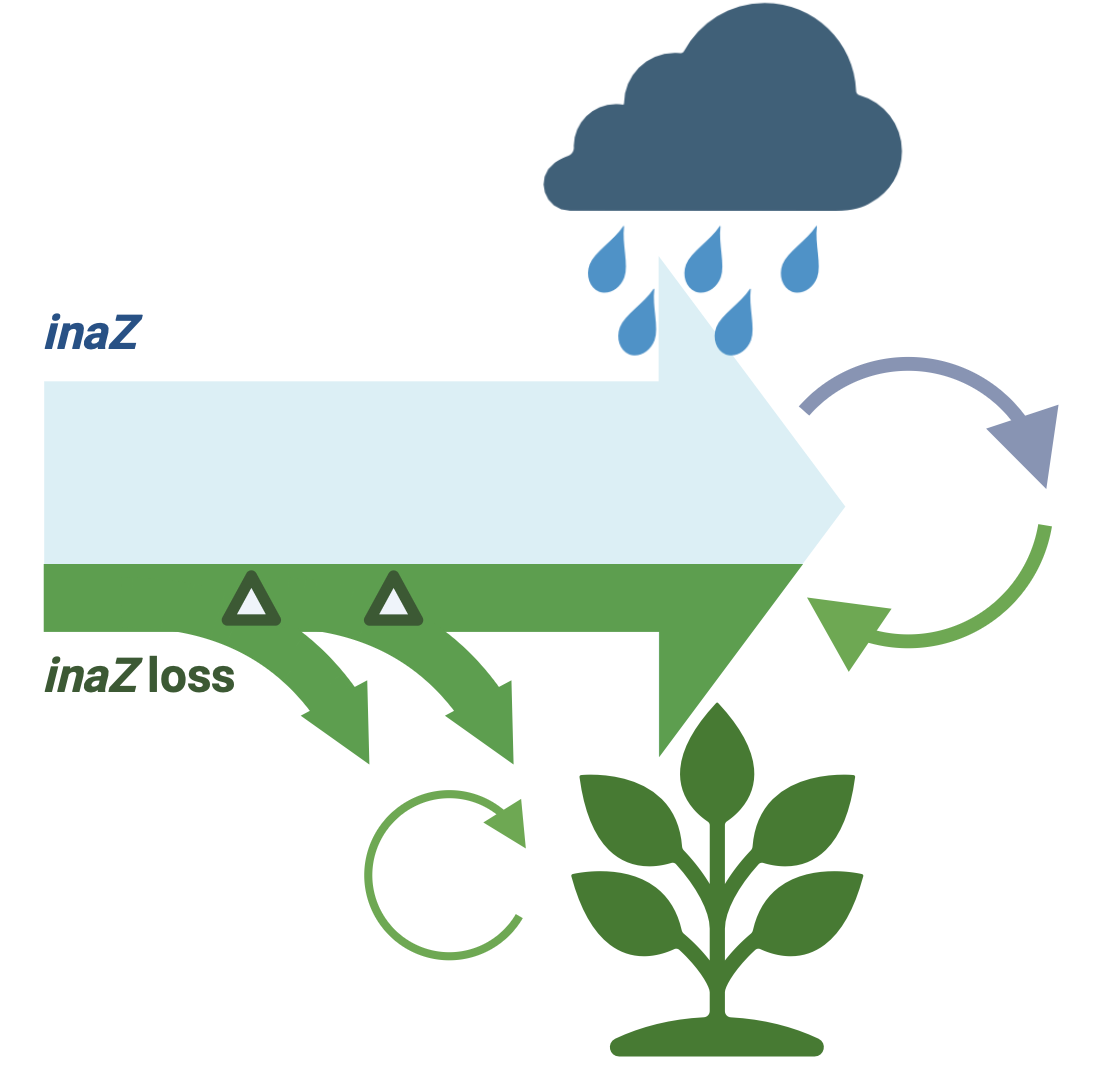

Fig. S7. Model for *inaZ* loss. Bacteria with *inaZ* play important roles in the water cycle. Bacteria that lose *inaZ* can no longer catalyze the formation of ice in clouds by freezing water and therefore cannot disperse via precipitation. The bacteria without InaZ then depend on distinct niches including host plant tissues for multiplication and dispersal.

Tables

Table S1. Metadata for *inaZ* analysis, whole genomes analyzed and mycotools databases.

Table S2. Hits for functional InaZ

**SI References**

1.	C. E. Morris, *et al.*, The life history of the plant pathogen Pseudomonas syringae is linked to the water cycle. *The ISME Journal* (2008). https://doi.org/10.1038/ismej.2007.113.
