## Supplementary material for "Loss of dispersal via ice nucleation activity constrains microbial evolution": BLASTData

inaZ orthologs figures


#### Table of contents

- 1 Species tree (1 genome per taxon)
  - 1.1 Without labels
- 2 Protein gene tree
- 3 Species tree vs. gene tree

### *inaZ* orthologs figures

Author

Jelmer Poelstra

Published

December 15, 2025


---

#### 1 Species tree (1 genome per taxon)

##### 1.1 Without labels

#### 2 Protein gene tree

#### 3 Species tree vs. gene tree
